# A tomato prolyl-4-hydroxylase causes relocation of abscission zone and alters abscission kinetics

**DOI:** 10.1101/2021.04.20.440677

**Authors:** Andreas Perrakis, Dusan Denic, Konstantinos N. Blazakis, Eleni Giannoutsou, Dimitrios Kaloudas, Craita E. Bita, Myrto Rizou, Afrodite Krokida, Mohamed Kouhen, Athina Lazaridou, Khansa Mekkaoui, Samia Belaidi, Zeina El Zein, Mohab Khalil, Lamia Ezzat, Noureldine Youssef, Maria Kosma, Anna G. González, Aline Monzer, Dimitra Papantoniou, Antri Varnava - Tello, Mondher Bouzayen, Ioannis-Dimosthenis S. Adamakis, Azeddine Driouich, Costas G. Billiaderis, Nicolas Kalogerakis, Panagiotis Kalaitzis

## Abstract

The detachment of organs is controlled by highly regulated molecular mechanisms. The position of the tomato abscission zone (AZ) is defined by the ratio of the proximal to distal part of the pedicel. In this study, the ratio was altered due to a shift in the position of the AZ which was attributed to shorter and longer pedicels of SlP4H3 RNAi and OEX lines due to changes on cell division and expansion in AZ and distal part. This might be associated with LM2- and JIM8-AGPs which increased in OEX and decreased in RNAi lines throughout the pedicel. The JIM13 AGPs were downregulated in the flower AZ of OEX lines, pointing to a role on abscission regulation. In addition, Co-IP in flower AZ with SlP4H3-GFP fusion proteins showed interaction with LM2-, JIM13- and JIM8-epitopes suggesting proline hydroxylation by SlP4H3. The lower content of methyl-esterified HGs and higher of demethyl-esterified HGs in the AZs of RNAi lines might be responsible for increased rigidity of the AZ cell walls, accounting for the higher force required for AZ tissue detachment to occur. Moreover, ethylene-induced flower abscission was accelerated in the RNAi lines and delayed in OEX lines, while exactly the opposite response was observed in the red ripe fruit AZs. This was partly attributed to alterations in the expression of cell wall hydrolases. Overall, these results indicate that P4Hs might regulate molecular and structural features of cell walls in the AZ as well as abscission progression by regulating the structure and function of AGPs.

**Sentence Summary:** Alterations on the expression of tomato prolyl 4 hydroxylase 3 shifts the position of the pedicel abscission zone and alters ethylene induced abscission progression according to the developmental context of the pedicel.

## Introduction

Abscission is the process of organ separation, a highly coordinated biological programme regulated predominantly by ethylene and auxin, while hormones such as giberrelins, cytokinins, abscisic acid and jasmonic acid also participate in the timing of abscission initiation (Tucker and Kim, 2015; Tranbarger and Tadeo, 2020; Sundaresan et al., 2016). The primary abscission model for plant species are Arabidopsis, tomato and soybean for flower organ, flower and fruit pedicel and leaf abscission, respectively.

In tomato, sensitivity to ethylene is considered essential for abscission competence and is acquired after auxin depletion (Meir et al., 2015; Guan et al., 2014). Abscission cell wall specific hydrolases such as polygalacturonases and cellulases are required for tomato abscission progression (Meir et al., 2010; Kalaitzis et al., 1995; Jiang et al., 2008).

The tomato flower abscission zone (AZ) has the shape of a distinct swollen node localized in the middle of the pedicel and is comprised of small, cytoplasmically dense cells of arrested differentiation (Tabuchi et al., 2001; Liu et al., 2014). The differentiation of the tomato pedicel AZ is regulated primarily by the *jointless* (*j*) mutant. Suppression of the SEPALLATA MADS-box TF (SlMBP21) was shown to abolish the development of the flower AZ, while overexpression resulted in shorter proximal pedicel (Liu et al., 2014).

Prolyl 4 hydroxylases (P4Hs) catalyze the hydroxylation of proline residues in Hydroxyproline Rich Glycoproteins (HRGPs) as the first step for subsequent O-glycosylation. Their physiological significance was demonstrated with three Arabidopsis P4Hs which affected root hair elongation by causing defects in the proper glycosylation of extensins (Velasquez et al., 2015). In addition, a cell surface Arabinogalactan peptide, AGP21, was shown to play a role in the root hair cell differentiation in Arabidopsis (Borassi et al., 2020). The post-translational O-glycosylation of AGP21 was a pre-requisite for correct targeting and functional conformation indicating the importance of proline hydroxylation (Borassi et al., 2020).

The peptide Inflorescence Deficient in Abscission (IDA) play the role of an abscission signal in Arabidopsis, undergoing proline hydroxylation, which was considered important for higher activity (Butenko et al., 2014). Recently, an IDA-like gene, (SlIDL6) was shown to promoter tomato flower abscission under low light conditions by upregulating AZ cell wall hydrolases (Li et al., 2021).

An additional peptide hormone, phytosulfokine (PSK), was found to regulate tomato flower abscission under drought conditions (Reichardt et al., 2020).The production of the active peptide depends on the highly regulated activity of a subtilisin-like protease, phytaspase 2 (SlPHYT2) (Reichardt et al., 2020). Overexpression of SlPHYT2 promoted abscission under drought stress while silencing delayed abscission (Reichardt et al., 2020).

In tomato, VIGS-induced silencing of three tomato P4Hs, SlP4H1, SlP4H7 and SlP4H9, resulted in longer shoots and roots and larger leaves due to increased cell division and expansion which was associated with a decrease in JIM8 AGPs and JIM11 extensins (Fragostefanakis et al., 2014). Moreover, RNAi-induced suppression of a prolyl 4 hydroxylase, SlP4H3, caused a delay in the abscission progression of overripe tomato fruits which was accompanied by significant down regulation of AZ cell wall hydrolases such as TAPG2, TAPG4, and CEL2 (Perrakis et al., 2019). The presence of Arabinogalactan proteins (AGPs) in Arabidopsis abscission was initially identified in Arabidopsis floral AZs (Stenvik et al., 2006). AGPs epitopes were also present in floral AZs indicating an increase in the AGPs (Stenvik et al., 2006). However, a decrease in the AGPs and RG-I domains (Rhamnogalacturonan-I) was observed in olive ripe fruit AZs during the separation process, accompanied by an increase in the de-esterified HGs (Homogalacturonan) (Parra et al., 2020).

Petal abscission in rose (*Rosa hybrida*) was regulated by two Ethylene Response Factors (ERFs) through the modulation of a β-galactosidase (RhBGLA1) expression (Gao et al., 2019). Silencing of the two ERFs accelerated petal abscission and induced a reduction of pectin epitopes (Gao et al., 2019). In tomato flower AZs, an increase in LM5-bound galactan and LM6-bound arabinan was observed while in fruit AZs no deposition of these polysaccharides was found (Iwai et al., 2013). In this report, the levels of SlP4H3 expression caused alterations on either the content or structure of AGPs and pectins in the flower AZs and pedicel, causing changes in cell division and cell expansion as well as in expression patterns of AZ cell wall hydrolases. As a result, at a structural level, the position of the AZ was shifted closely and distantly to subtending organ due to shorter and longer distal pedicel, respectively. At the developmental level, the abscission rate of open flower and red ripe fruit was either accelerated or delayed depending on the expression levels of SlP4H3.

## Results

### Overexpression and RNAi SlP4H3 lines exhibit pedicel abscission acceleration and delay under ethylene

Silencing of three SlP4H3 RNAi lines delayed natural fruit abscission 90 days after the breaker stage (Perrakis et al., 2019). To further investigate the involvement of SlP4H3 in abscission, a construct comprising the 35S promoter directing the expression of this cDNA was transformed into microtome tomato plants by using *A. tumefaciens* mediated transformation. Seven independent microtome overexpression lines were produced and five were crossed with the wild type *Ailsa Craig* tomato plants. Two homozygous overexpression lines, SlP4H3 OEX#1 and SlP4H3 OEX#2, were selected for further characterization after validating higher transcript abundance ranging at the levels of 19-to 4-fold in various tissues such as, leaf, stem, root and fruits at growth and breaker stages (**Figure S1**).

The Arabidopsis P4H2, P4H5 and P4H13 polypeptides were comprising of signal peptides and shown to be localized in the ER and Golgi network (Velasquez et al., 2015). However, no signal peptide was present in the SlP4H3 peptide, only a transmembrane domain. Stable, independent tomato transformant lines were produced carrying a construct of the P4H3 cDNA with a GFP tag at the C-terminus under the control of the 35S promoter. The construct was localized in the ER membrane and nuclear envelope in leaf and root cells of young seedlings (**Figure 1**).

**Figure 1.**
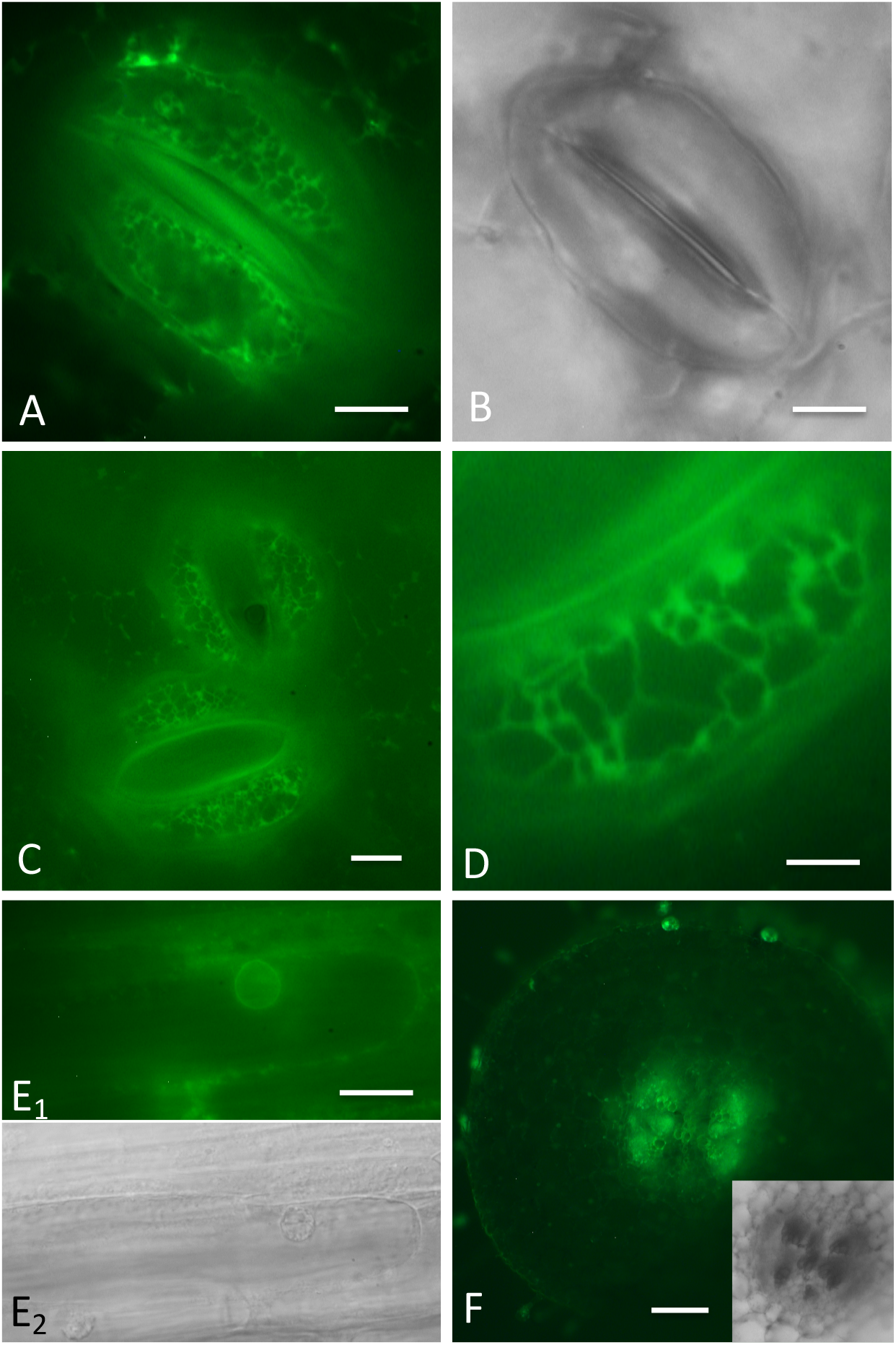
Expression of P4H3 cDNA construct with a C-terminus GFP tag under the control of the 35S promoter in stable tomato independent transformant line. The GFP protein was localized in the ER membrane and the nuclear envelope of the leaf cells **(A-D)** and at the nuclear envelope of the root cells **(E1-E2)**. At the transverse section of the shoot, the GFP protein is located at the phloem elements of the vascular tissue **(F) ? –D**: Leaves paradermal section **E:** Root epidermal cells **F**: Shoot transverse section. Scale bars: A-C: 10 μm, D: 5 μm, E: 10 μm, F: 100 μm.

The expression levels of SlP4H3 in the RNAi and OEX lines was determined in an ethylene-induced flower and red ripe fruit pedicel abscission time course (**Figure 2C, D**). Higher expression of SlP4H3 ranging between 6-to 4-fold in OEX lines were observed throughout the entire time course (**Figure 2C, D**) while in the RNAi lines significant suppression of expression was recorded (**Figure 2C, D**).

**Figure 2.**
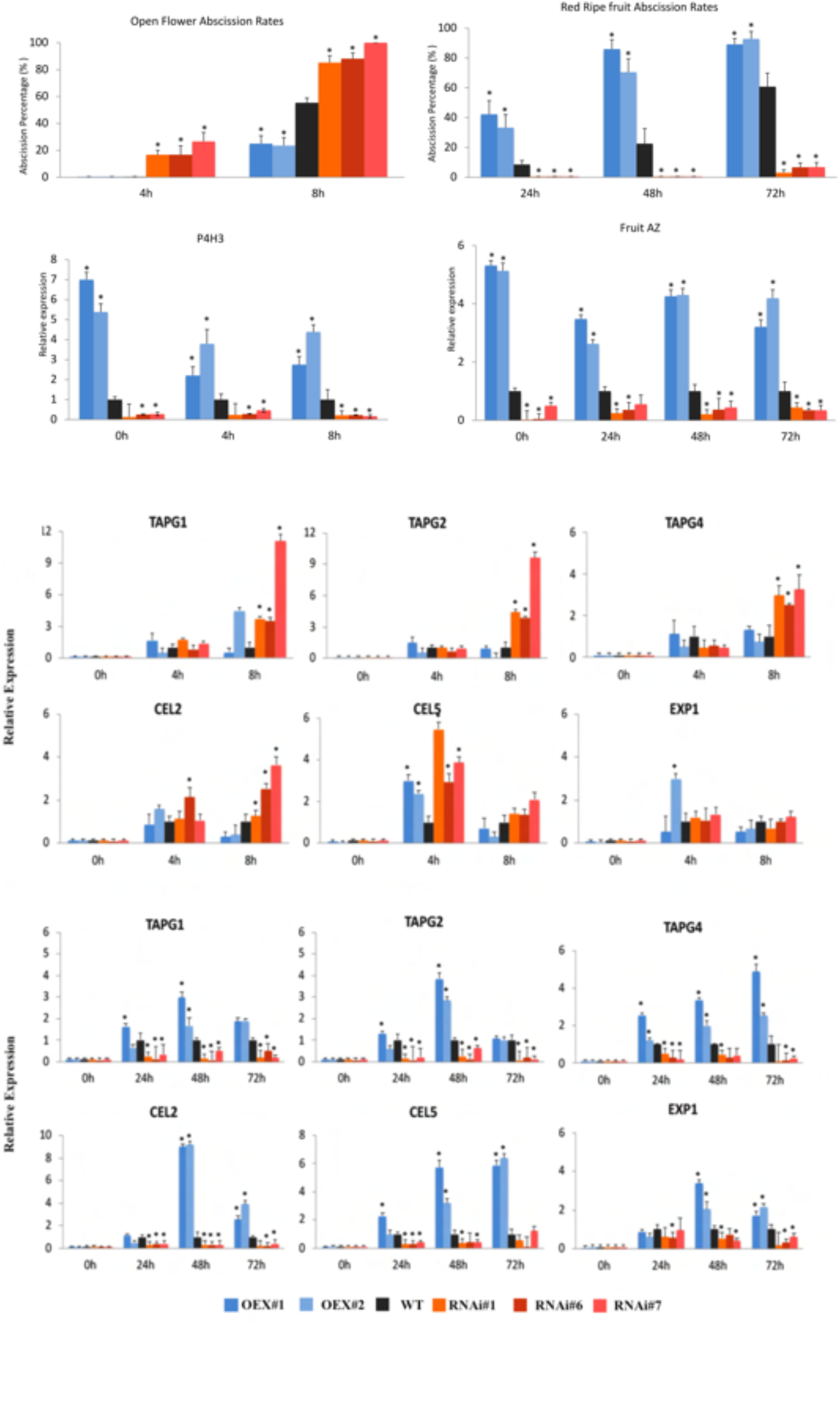
Time course of ethylene-induced abscission rates of flower and fruit AZs and expression of SlP4H3, SlTAPG1, SlTAPG2, SlTAPG4, SlCEL2, SlCEL5 and SlEXP1 (expansin) in RNAi #1, #6, #7 and OEX #1, #2 lines and wild type. **A**. Time course of abscission rates in open flower AZs for 4 and 8 hour. **B**. Time course of abscission rates in red ripe fruit AZs for 24, 48 and 72 hour. **C**. Relative expression of SlP4H3 in red ripe fruit AZs of RNAi, OEX lines and wild type. **D**. Relative expression of SlP4H3 in open flower AZs of RNAi, OEX lines and wild type. **E**. Relative expression of SlTAPG1, SlTAPG2, SlTAPG4, SlCEL2, SlCEL5 and SlEXP1 in AZs of open flowers after 4 and 8 hour of ethylene in RNAi, OEX lines and wild type. **F**. Relative expression of SlTAPG1, SlTAPG2, SlTAPG4, SlCEL2, SlCEL5 and SlEXP1 in AZs of red ripe fruit after 24, 48 and 72 hour in RNAi, OEX lines and wild type. The relative expression was calculated according to the comparative Ct method by using actin as internal standard. The asterisk indicates statistically significant differences. The subtending flowers and fruits were removed from the explants and then were treated with 100 ppm of ethylene while the 0 hour refer to the un-treated control.

Abscission explants after removal of the subtending organs at the stage of open flower and red ripe fruit were treated with ethylene for 8 and 72 hours, respectively (**Figure 2A, B**). At the flower stage, the three RNAi lines, #1, #6 and #7 showed a significant acceleration of pedicel abscission compared to wild type while the two OEX lines, #1 and #2 exhibited a significant delay (**Figure 2A**). The abscission was higher than 80% for the three lines after 8 hours compared to 60% and approximately 20% rate of the wild type and the two OEX lines, respectively (**Figure 2A**).

Remarkably, exactly the opposite trend in abscission was observed for the RNAi compared to the OEX lines at the red ripe fruit stage (**Figure 2B**). The RNAi lines showed a considerable delay in abscission, at levels of 10% after 72 hours, while the OEX lines exhibited an acceleration in abscission which was higher than 80%, compared to a 60% abscission found for the wild type (**Figure 2B**). This fruit abscission delay in the RNAi lines is in accordance with the delay in the natural over ripe fruit abscission 90 days after the breaker stage (Perrakis et al., 2019).

### Flower and fruit abscission progression in relation to expression patterns of cell wall hydrolases

The expression of abscission cell wall hydrolases was determined in order to further investigate the underlying physiological mechanisms for the observed opposite trends in abscission progression.

The TAPG1, TAPG2, TAPG4 and CEL2 were significantly up-regulated in the flower AZs of RNAi lines compared to wild type after 8 hour of ethylene (**Figure 3E**). Instead, no significant variation in the expression of these genes was observed in the OEX lines (**Figure 3E**). Only a 4-fold increase of TAPG1 in OEX line #2 was observed after 8 hours of ethylene treatment (**Figure 3E**). The CEL5 mRNA was up-regulated in RNAi and OEX AZs after 4 hours, while no significant differences in expression were observed after 8 hours of ethylene (**Figure 3E**). No alterations in the expression of Expansin 1 (EXP1) were observed neither in RNAi nor in OEX lines except an increase in OEX line #2 after 4 hours of ethylene (**Figure 3E**). These results indicate that the higher abscission in RNAi lines might be partly attributed to the up-regulation of cell wall hydrolases after 8 hours of ethylene (**Figure 3E**).

**Figure 3.**
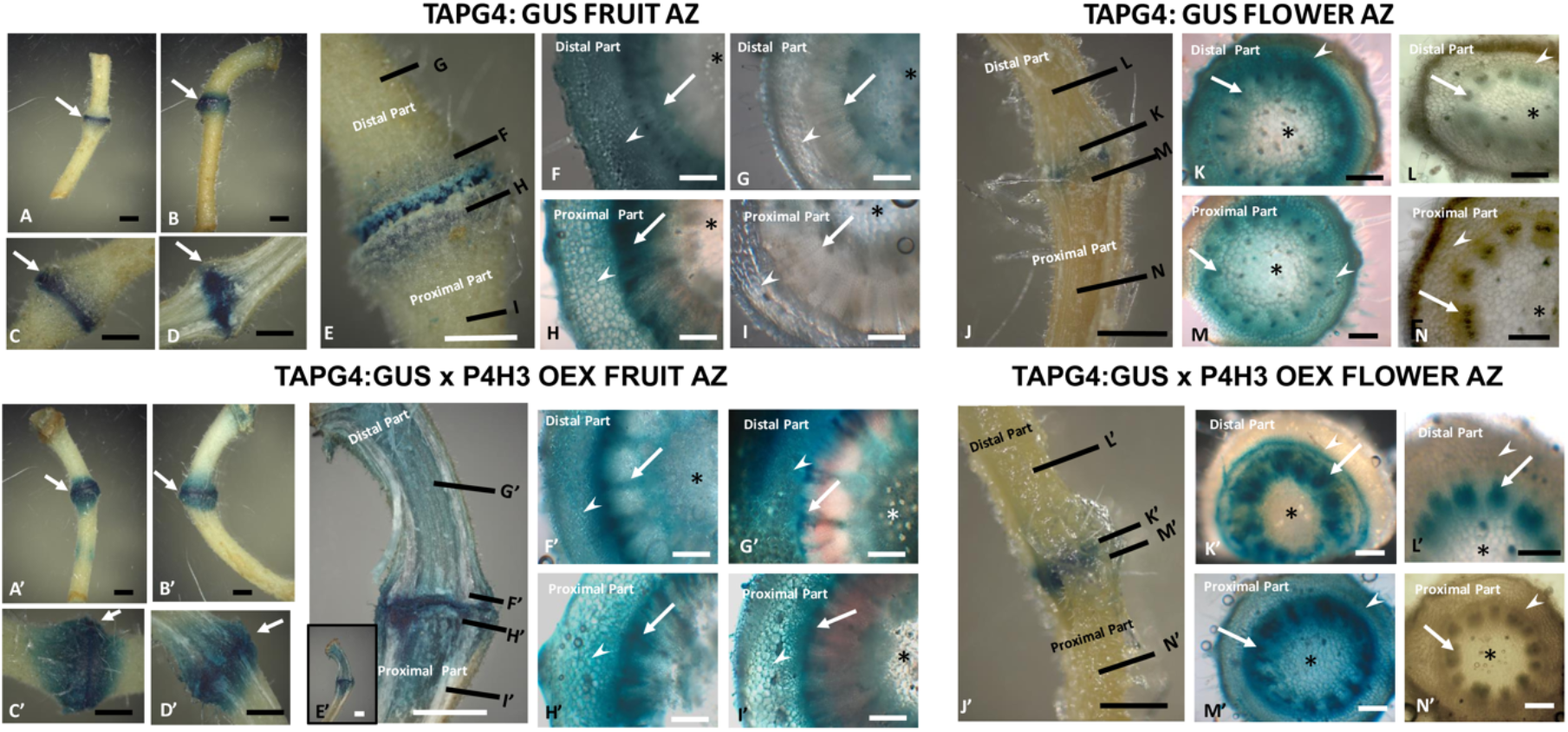
TAPG4:GUS fruit abscission layers in A-I and TAPG4: GUS x P4H3 OEX in A’-I’. TAPG4: GUS flower abscission layers in J-N and TAPG4: GUS x P4H3 OEX in J’-N’. A-C and A’-C’: Non dissected fruit abscission zone pedicels, while D, D’: dissected fruit abscission zone pedicel. The arrows point to the fruit abscission layers stained in blue. In E a non dissected fruit AZ while in E’ a longitudinally dissected fruit AZ. Inset in E’: the whole fruit pedicel, shown in magnification in E’. In both pictures, the black lines mark the planes where the transverse sections shown in F-I and F’ –I’ were taken. F-I and F’-I’: transverse section at different sites marked by the respective letters in E and E’. The arrows point to the xylem of the vascular bundles, the asterisks mark the pith, while the arrowheads point to the cortex cells. In J and J’ a non dissected flower AZ. In both pictures, the black lines mark the planes where the transverse sections shown in K-N and K’ –N’ were taken. K-N and K’-N’: transverse section at different sites marked by the respective letters in J and J’. The arrows point to the xylem of the vascular bundles, the asterisks mark the pith, while the arrowheads point to the cortex cells. Scale bars in A-D and A’-D’: 600μm, E and E’: 300 μm scale bars in F-I and F’-I’: 100μm, Scale bars in J-J’: 400μm and scale bars in K-N and K’-N’: 100μm.

In the fruit AZs, all three RNAi lines exhibited a remarkable decrease in expression for TAPG1, TAPG2, TAPG4, CEL2, CEL5 and EXP1 in the ethylene time course (**Figure 3F**). However, in the OEX lines, the TAPG1 and EXP1 expression peaked after 48 hours in line #1 while in line #2 the mRNA levels increased after 48 and 72 hours (**Figure 3F**). The TAPG2 and CEL2 showed an upregulation at 48 hours by approximately 3- and 9-fold, respectively while the TAPG4 and CEL5 exhibited a gradual increase in expression throughout the time course (**Figure 3F**). These results suggest that the higher abscission rates in OEX lines might be partly justified by the up-regulation of cell wall hydrolases.

A stable transformant line expressing the GUS reporter cDNA under the control of the TAPG4 promoter (Hong et al., 2000) was crossed with the SlP4H3 OEX #1 to further investigate the temporal and spatial expression of TAPG4 in the OEX background (**Figure 3**). Flower and red ripe fruit AZ of double homozygous explants were treated with ethylene for 8 and 72 hours, respectively (**Figure 3**). TAPG4:GUS staining indicated overall higher expression in the ethylene treated fruit AZs of OEX#1 line validating the qPCR data (**Figure 3**). The GUS staining in the flower AZs after 8 hour of ethylene was similar in OEX #1 and wild type (**Figure 3**).

The GUS staining was more intense in ethylene-treated OEX fruit AZ layers compared to wild type (**Figure 3A’-E’, 3A-E**) while the GUS levels remained high at the entire distal part (**Figure 3E’**) and at a large part of the proximal (**Figure 3E’)**. At the wild type cross sections on both sides of the AZ, GUS staining was detected at the cortex parenchyma cells, vascular tissue and the pith (**Figure 3F, 4H)**. At cross sections distant from the AZ, GUS staining is fading showing only a weak signal at the protoxylem vessels and the pith at the distal (**Figure 3G**); and proximal part of the pedicel (**Figure 3I**). At the OEX line, GUS staining is intense at all tissues on both sides of the AZ (**Figure 3F’**,**4H’)** and remained at very high levels at cross sections away from the AZ at the distal (**Figure 3G’**) and proximal part (**Figure 3I’**). The staining is obvious at the cortex parenchyma cells (arrowheads in **Figure 3G’** and **Figure 3I’**) and at the newly produced xylem vessels near cambium (arrows in **Figure 3G** and **Figure 3I’**).

GUS staining was observed at all tissues near the AZ (**Figure 3K** and **Figure 3M**), both at the proximal and distal part of TAPG4: GUS flower line. At cross sections distant from the AZ, faint GUS staining was only visible at the protoxylem vessels at the distal part (**Figure 3L**). No GUS staining was observed at distant from the AZ cross sections of the proximal pedicel (**Figure 3N**). In the OEX lines, the GUS signal was more intense at the distal part compared to wild type. More precisely, GUS staining was observed in the vascular tissue, but also at the parenchyma cells that encompass the ring of the vascular bundles, as has already been described by Hong et al. (2000) (arrowhead in **Figure 3K’**). At the proximal pedicel near the AZ, the GUS staining was intense at all the tissues examined (**Figure 3M’**). At a distance from the AZ, GUS staining was not observed at the proximal pedicel (**Figure 3N’**), whereas was still detected at the xylem vessels of the distal pedicel (**Figure 3L’**). These results indicate that overexpression of SlP4H3 induces ectopic expression of TAPG4 in the entire distal pedicel but not in the proximal part.

### Shift in the position of the AZ in RNAi and OEX lines

The distal part of the pedicel was shorter in RNAi and longer in OEX lines compared to the wild type (**Figure 4A, D, Figure S2**). Similar results were obtained with the distal part length in the red ripe fruits (**Figure S2A**). This shift in the position of the AZ is nicely reflected in the ratio of proximal to distal pedicel length (**Figure 4F**) which determines the position of the AZ within the pedicel. Higher ratio means higher proximity of the AZ to the flower while lower ratio indicates lower proximity to the flower (**Figure 4F**). Moreover, the average fruit AZ diameter in OEX lines was smaller, compared to the wild type and RNAi (**Figure S2A**).

**Figure 4.**
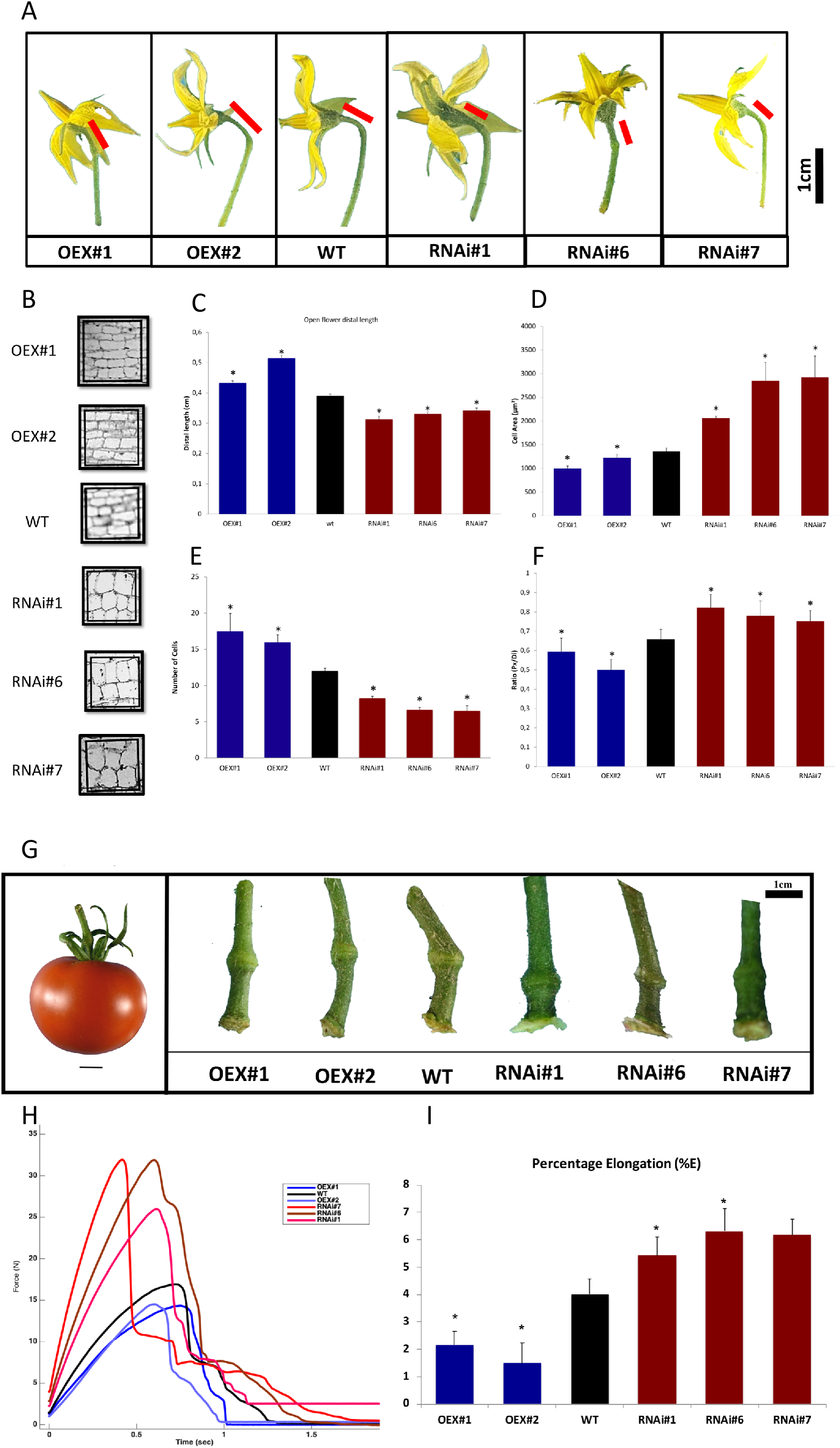
Flower Pedicel characterization and fruit AZ detachment Force and Percentage Elongation (%E) in the RNAi #1, #6, #7 and OEX #1, #2 lines and wild type. **A**. Representative open flower pedicels of the OEX and RNAi lines and wild type. Red lines depict the length of the distal part of the pedicel. **B**. Longitudinal sections depicting indicative cell number and cell area of distal pedicel parenchymatic cells in a specific square area of 196 μm^2^. **C-F**. The length of the distal part of the pedicel (n=180 pedicels/line), the cell area and number of cells in a specific square area of the distal pedicel parenchymatic cells in OEX and RNAi lines(cell morphology measurements made on 9 images per line; three regions per AZ from three different plants of the same line). The ratio of the proximal pedicel length to distal pedicel length as indicator of the position of the abscission in OEX and RNAi lines (n=180 pedicels/line). **G**. Representative red ripe fruit pedicels of the OEX and RNAi lines and wild type. **H**. The Detachment Force is presented as the diagram of breakstrength force (Newton) required for detachment to occur in a particular period of time (seconds) in red ripe fruit AZ explants of RNAi #1, #6, #7 and OEX #1, #2 lines and wild type. The RNAi lines require higher detachment force in a shorter period of time compared to OEX lines which require lower force and longer time. The detachment force of wild type is placed in between the RNAi and OEX lines. **I**. The percentage elongation (%E) associates the deformation of the tissue before the failure stage. Higher %E was observed in the RNAi #1, #6, #7 lines compared to the OEX #1, #2 lines indicating higher adhesiveness. The AZ toughness and texture, adhesive tests were performed by using a texture analyzer (n=20 pedicels/line). The asterisks indicate statistically significant differences compared to wild type.

In the distal part of OEX lines, the number of parenchymatic cells is larger in a defined square area (**Figure 4B, E**). The average cell number is higher than 15, compared to 12 cells in wild type and less than 7 in RNAi lines (**Figure 4B, E**). The cell area is smaller in OEX lines and larger in RNAi lines, compared to the wild type (**Figure 4B, D**). These data imply alterations in both cell division and expansion.

The tomato flower AZ is comprised of 5-10 rows of small, dense, disk-shaped cells while the separation layer is usually composed of 1-5 cell layers (Roberts et al., 1984). Differences were observed in the number of the fruit AZ cell rows in the OEX lines, mainly within the cortex and the pith parenchymatic cells. The number of fruit AZ cell layers in OEX lines ranged at the level of 17-20 compared to 10 cell layers in RNAi and wild type (**Figure S2B, C)**. Moreover, OEX and RNAi AZ cells appeared more rectangular in comparison to wild type in which the circular, spherical morphology is dominant (**Figure S2B, C)**. The differences in cell shape can be easily deducted by the higher cell width, while no differences were observed in circularity (**Figure S2B, C, D**). These results indicate rather similar cell shape characteristics between RNAi and OEX lines.

### The fruit AZ of RNAi lines require higher force of detachment and percentage elongation (%E) for separation

Tensile tests were performed to assess the AZ toughness and deformation characteristics. Clearly, the Young’s modulus values (initial slopes of the force-deformation curves) were greater for the RNAi lines, as a group, compared to OEX lines. Two other characteristics were also determined: the detachment force, obtained as the maximum force versus time (deformation) diagram, which can reveal differences in mechanical behavior; (ii) percentage elongation (%E) which expresses the deformation of the AZ up to the failure point (**Figure 4H**). The break strength force required for abscission to occur in RNAi was significantly higher compared to the wild type (**Figure 4H**). A lower detachment force was recorded in the case of OEX lines (**Figure 4H**).

The %E in the OEX AZs was lower by 50%, compared to wild type, while for the three RNAi lines was higher by 32%, 55% and 54% (**Figure 4I**). In the RNAi pedicels, the higher %E is also reflected by the larger width of the force-deformation curves, compared to OEX and wild type, indicating greater extension upon tissue detachment (**Figure 4H**). The stress-deformation profiles suggest that the OEX pedicels resemble more the mechanical failure of brittle materials, whereas the RNAi pedicels exhibit a more ductile behaviour.

These mechanical deformation profiles clearly reveal distinct differences in the strength and %E of the AZ between the RNAi and OEX lines which might originate from varying structural, geometrical and morphological features of the fruit pedicel, AZ cell layers, cell size and probably cell wall and middle lamellae properties. The higher the breaking force and %E of the AZs, the more difficult is for abscission to occur either naturally or via ethylene treatment. These large deformation tensile tests were in agreement with the ethylene-induced red ripe fruit abscission rates observed between the RNAi and OEX lines.

### Immunolocalization of AGPs-bound epitopes showed higher expression in OEX lines and lower in the RNAi lines

The LM2, JIM13 and JIM8 antibodies were used in longitudinal sections of the flower pedicel AZ. Significantly lower abundance of the LM2 AGPs was detected in all three RNAi lines throughout the pedicel tissue and not only in the AZ cells compared to wild type (**Figure 5A-D**). On the contrary, the fluorescence signal intensity was much more intense in the AZ, proximal and distal cells of OEX lines compared to wild type (**Figure 5E, F**). These results indicate higher and lower expression of LM2 AGPs in OEX and RNAi lines throughout the pedicel, respectively (**Figure 5A-F**). The JIM8 AGPs showed lower signal intensity in the AZ, proximal and distal cells in all three RNAi lines compared to wild type (**Figure 5G-J**), while the OEX lines displayed fluorescent signal of higher intensity (**Figure 5K, L**, compare to **Figure 5G**). Immunolabeling by JIM13 antibody showed a similar epitope distribution between RNAi lines and wild type at the AZ cell area as well as in the proximal and distal area (**Figure 5M-P**). Observing the signal at the two OEX lines, indicates the following deposition pattern: the fluorescence signal in the AZ cells is less intense compared to the proximal and distal cells signal (**Figure 5Q, R**). The JIM13 AGPs showed an AZ-specific pattern of expression compared to LM2 and JIM8 AGPs (**Figure 5**). Overall, the expression of AGPs was higher in OEX lines and lower in RNAi lines either at the whole pedicel level or at the AZ-specific level, with the exemption of JIM13 expression in OEX lines (**Figure 5**).

**Figure 5.**
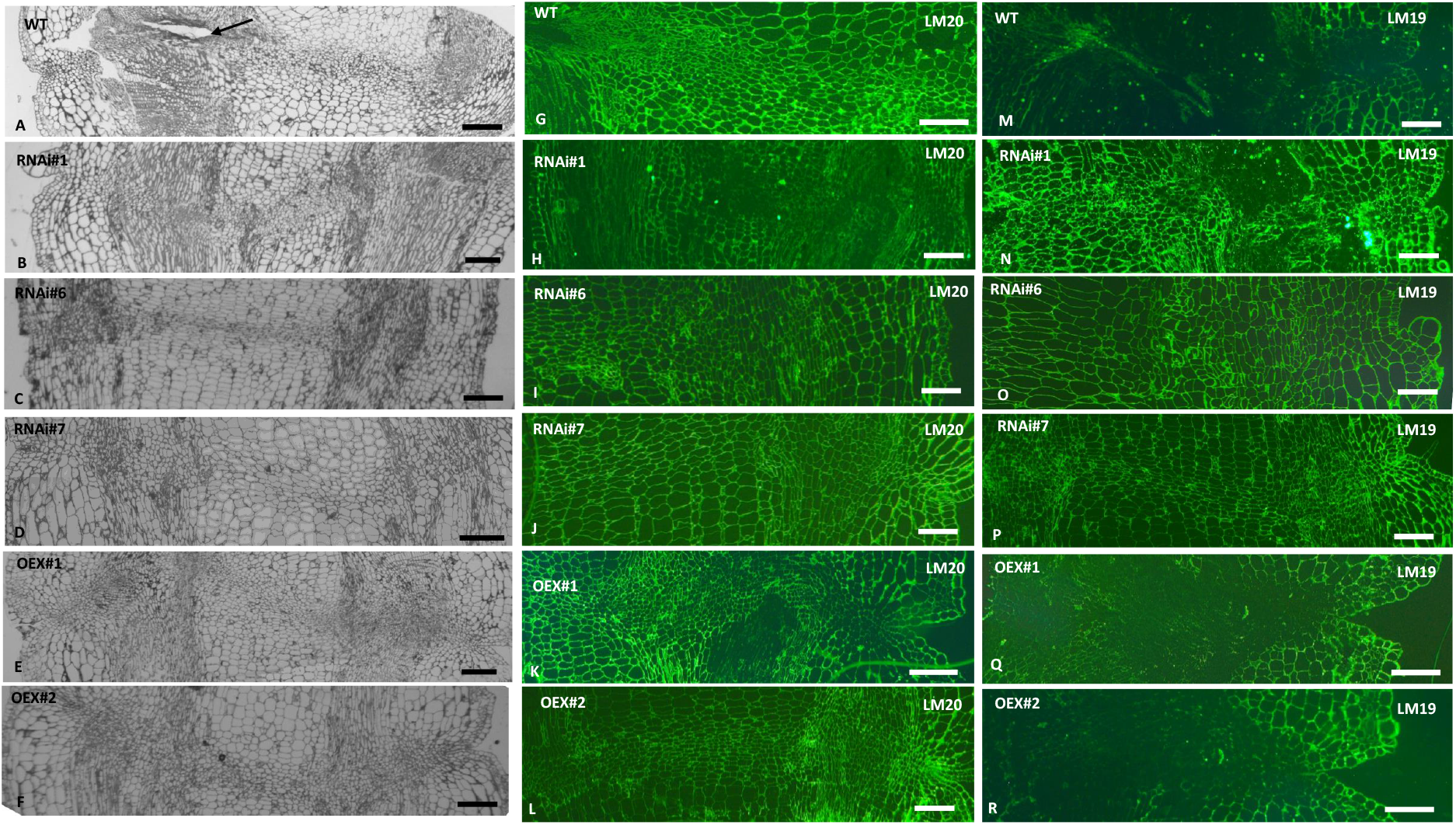
Immunolocalization of pectic epitopes in the flower AZ cells in OEX #1, #2 and RNAi #1, #6, #7 lines and wild type. (A-F) Longitudinal sections of flower AZ were stained with toluidine to determine the morphological characteristics of the cells at the AZ. (A)Wild type, (B)RNAi #1, (C)RNAi #6, (D)RNAi #7, (E)OEX#1, (F)OEX#2. (G-L) Longitudinal sections of flower AZ where immunolocalization using LM20 antibody was performed. (G)Wild type, (H)RNAi #1, (I)RNAi #6, (J)RNAi #7, (K)OEX#1, (L)OEX#2. (M-R) Longitudinal sections of flower AZ where immunolocalization using LM19 antibody was performed. (M)Wild type, (N)RNAi #1, (O)RNAi #6, (P)RNAi #7, (Q)OEX#1, (R)OEX#2. Scale bars =100μ?.

### Differential immunolocalization of methyl-esterified and demethyl-esterified homogalacturonans (HGs) in the AZ of RNAi and OEX lines

Monoclonal antibodies were used to stain longitudinal sections of the AZs in order to determine the distribution of pectic polysaccharides. The LM2 antibody binds to highly methylesterified homogalacturonans whereas LM19 binds to partially demethyl-esterified and non-methylesterified homogalacturonans. The methylesterified homogalacturonans are highly abundant y at the wild type AZ and less abundant in the distal and proximal part (**Figure 6G**). In the RNAi#1, clearly lower intensity was observed in AZ cells compared to distal and proximal regions (**Figure 6H**), while in the case of RNAi#6 and RNAi#7, the signal was of lower intensity across the pedicel tissue including the AZ compared to wild type (**Figure 6I, J**). The OEX lines displayed intense signal at the AZ area and at the surrounding tissues, with a similar to the wild type intensity (**Figure 6K, L**). These results indicate lower methyl-esterified homogalacturonans in the flower AZ cells of RNAi lines compared to wild type and OEX lines.

**Figure 6.**
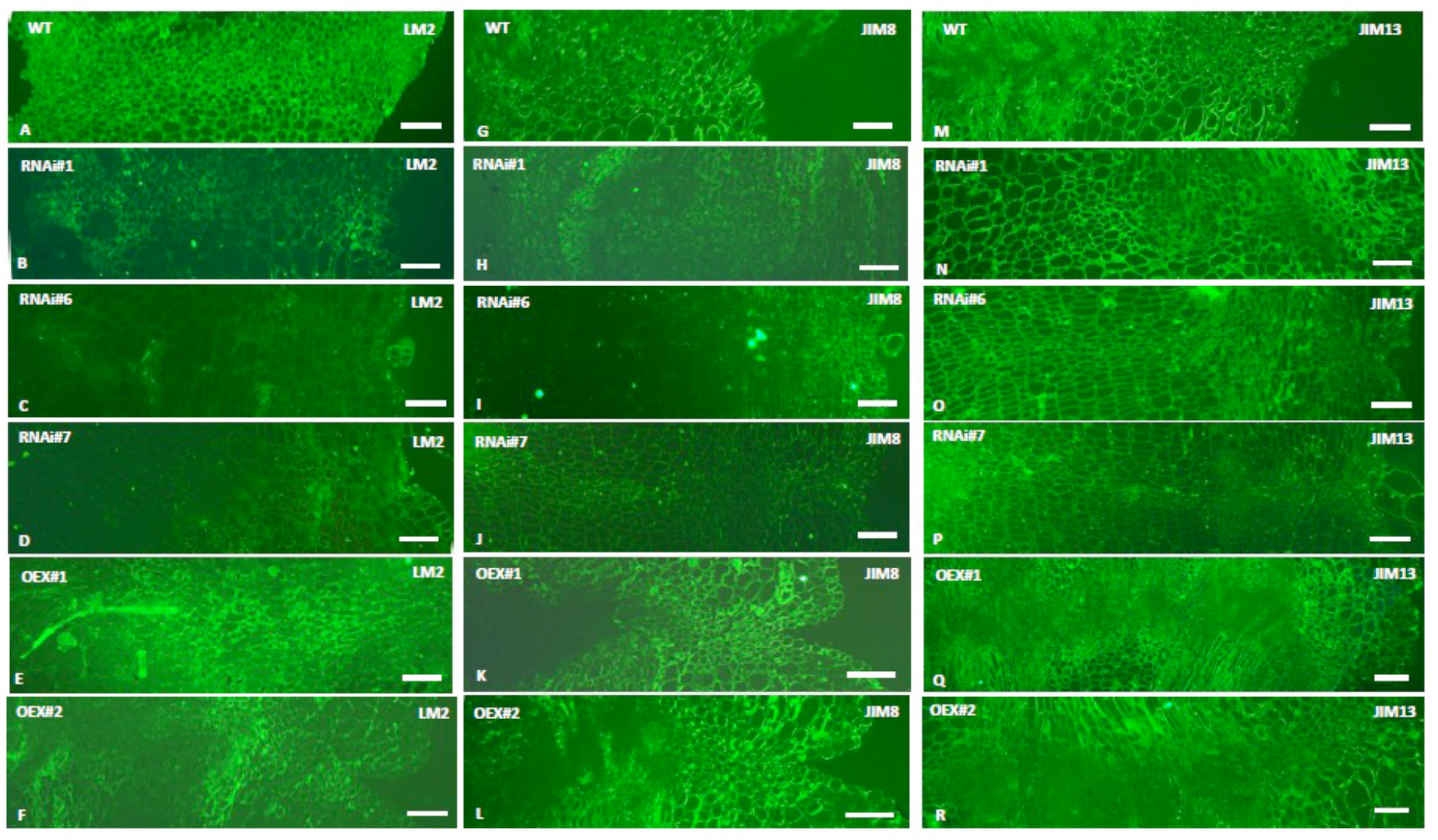
Immunolocalization of AGPs-bound epitopes in the flower AZ cells in OEX #1, #2 and RNAi #1, #6, #7 lines and wild type. (A-F) Longitudinal sections of flower AZ where immunolocalization using LM2 antibody was performed (A)Wild type, (B)RNAi #1, (C)RNAi #6, (D)RNAi #7, (E)OEX#1, (F)OEX#2. (G-L) Longitudinal sections of flower AZ where immunolocalization using JIM8 antibody was performed. (G)Wild type, (H)RNAi #1, (I)RNAi #6, (J)RNAi #7, (K)OEX#1, (L)OEX#2. (M-R) Longitudinal sections of flower AZ where immunolocalization using JIM13 antibody was performed. (M)Wild type, (N)RNAi #1, (O)RNAi #6, (P)RNAi #7, (Q)OEX#1, (R)OEX#2. Scale bars =100μ?.

Lower abundance of demethyl-esterified homogalacturonans were detected in wild type AZ cells compared to distal and proximal (**Figure 6M**) while, remarkably, in all RNAi lines, the LM19 fluorescent signal was intense, specifically in the AZ cells of RNAi #1 line (**Figure 6N-P**). Surprisingly, the OEX AZ cells did not display any signal at all whereas the distal and proximal cells displayed intense signal (**Figure 6Q, R**). Immunodetection with LM19 showed a specificity of this epitope only in the AZ of RNAi and not in wild type and OEX. These data imply that only the RNAi flower AZ cell walls contained demethyl-esterified homogalacturonans. Such a compositional difference for AZ pectins might have an impact on the ethylene-induced flower abscission.

### Interaction of SlP4H3 with AGPs-bound epitopes

To further investigate the interaction of SlP4H3 with AGPs in abscission, two constructs comprising the 35S promoter directing the expression of SlP4H3 cDNA with N- and C-terminal GFP fusions were stably transformed into Ailsa Craig and microtome tomato plants, respectively. The C-GFP microtome independent stable transformant lines were then crossed with Ailsa Craig. The GFP trap system was used to purify co-precipitated proteins from flower AZs of P4H3-GFP homozygous plants and then western blot analysis with the LM2, JIM13 and JIM8 antibodies was performed (**Figure 7**). The three AGP-bound epitopes were co-precipitated with the GFP antibody (beads fraction) indicating interaction with the SlP4H3. AGPs epitopes were also detected in the elution fraction (**Figure 7**). This suggest that SlP4H3 may interact with several members of the tomato AGPs family. The SlP4H3-GFP fusion protein of 61 KDa was detected in the co-precipated proteins in both N- and C-GFP lines but not in the wild type (**Figure 7**).

**Figure 7.**
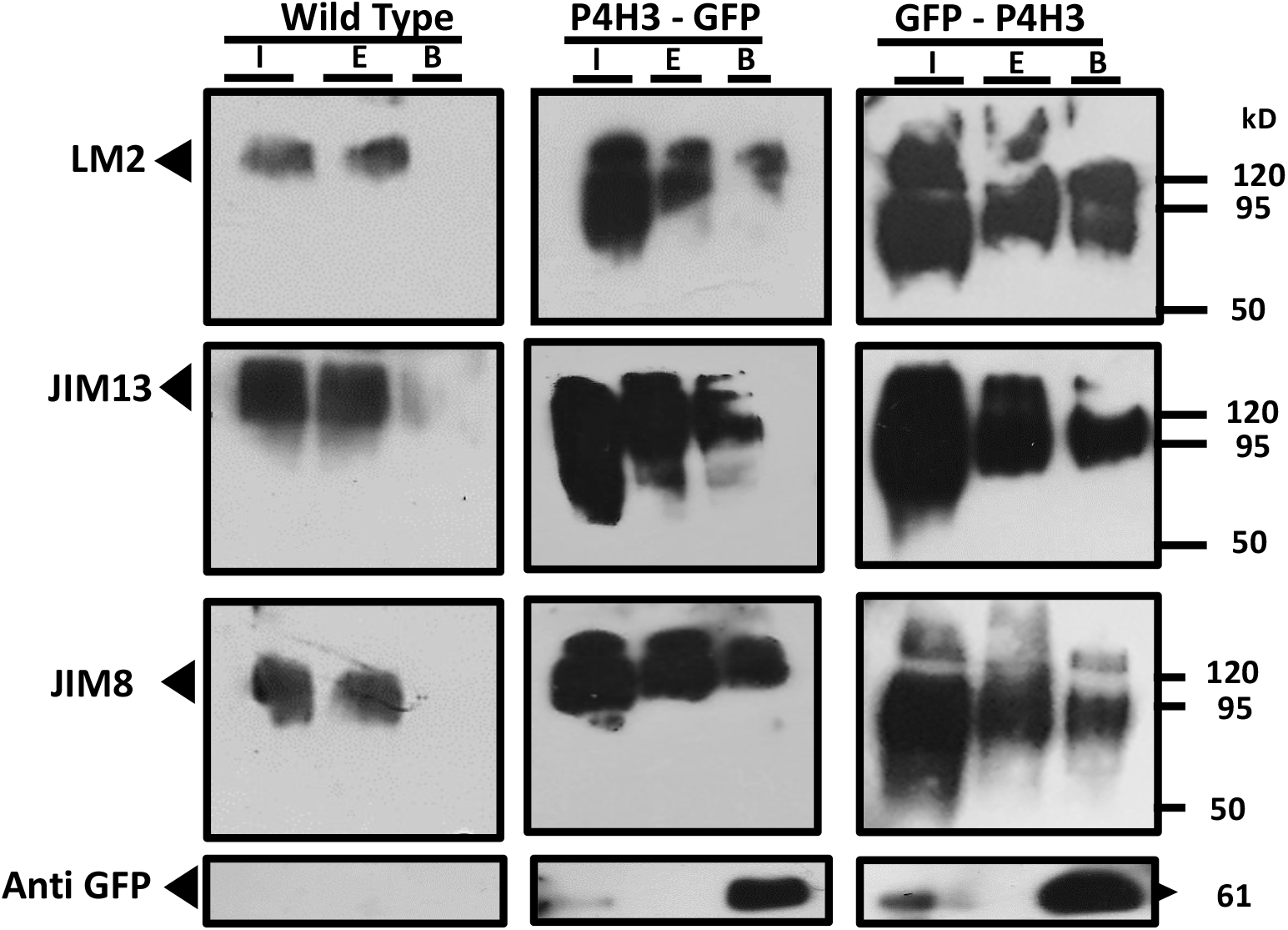
The SlP4H3 cDNA was transformed to produce overexpression N-terminal (GFP-P4H3) and C-terminal (P4H3-GFP) GFP stable tomato transformants to be used for Immunoprecipitation with GFP-trap. Co-immunoprecipitated protein complexes were analyzed by western blot analysis using the LM2, JIM13 and JIM8 antibodies. Flower AZs total protein extracts were used as input (I) for Co-IP and fractionated in SDS-PAGE with the elution (E) proteins and the precipitated by the GFP beads (B) material. Wild Type total protein extract was also used in a similar way as negative control. The N- and C-GFP fusion proteins of 61 KDa were detected by using the GFP antibody.

## Discussion

Suppression and overexpression of tomato SlP4H3 caused pleiotropic phenotypes in the AZs morphology, the pedicel length and the flower and fruit abscission programme. The abscission specific phenotypes are depicted in a graphic illustration of the molecular changes in the SlP4H3 and AGPs epitopes which cause changes in the morphology of AZ and pedicel as well as in the expression of cell wall hydrolases leading to differential abscission (**Figure 8**). Suppression of SlP4H3 shifted the position of the AZ closer to the subtending organ, while overexpression shifted the AZ further away (**Figure 8**). Moreover, ethylene-induced flower abscission was accelerated in RNAi lines and delayed in OEX lines (**Figure 8**), whereas exactly the opposite response was observed in the red ripe fruit AZs; ethylene accelerated and delayed abscission in OEX and RNAi lines, respectively. The AZ in a different developmental context responded in exactly the opposite way under ethylene treatment.

**Figure 8.**
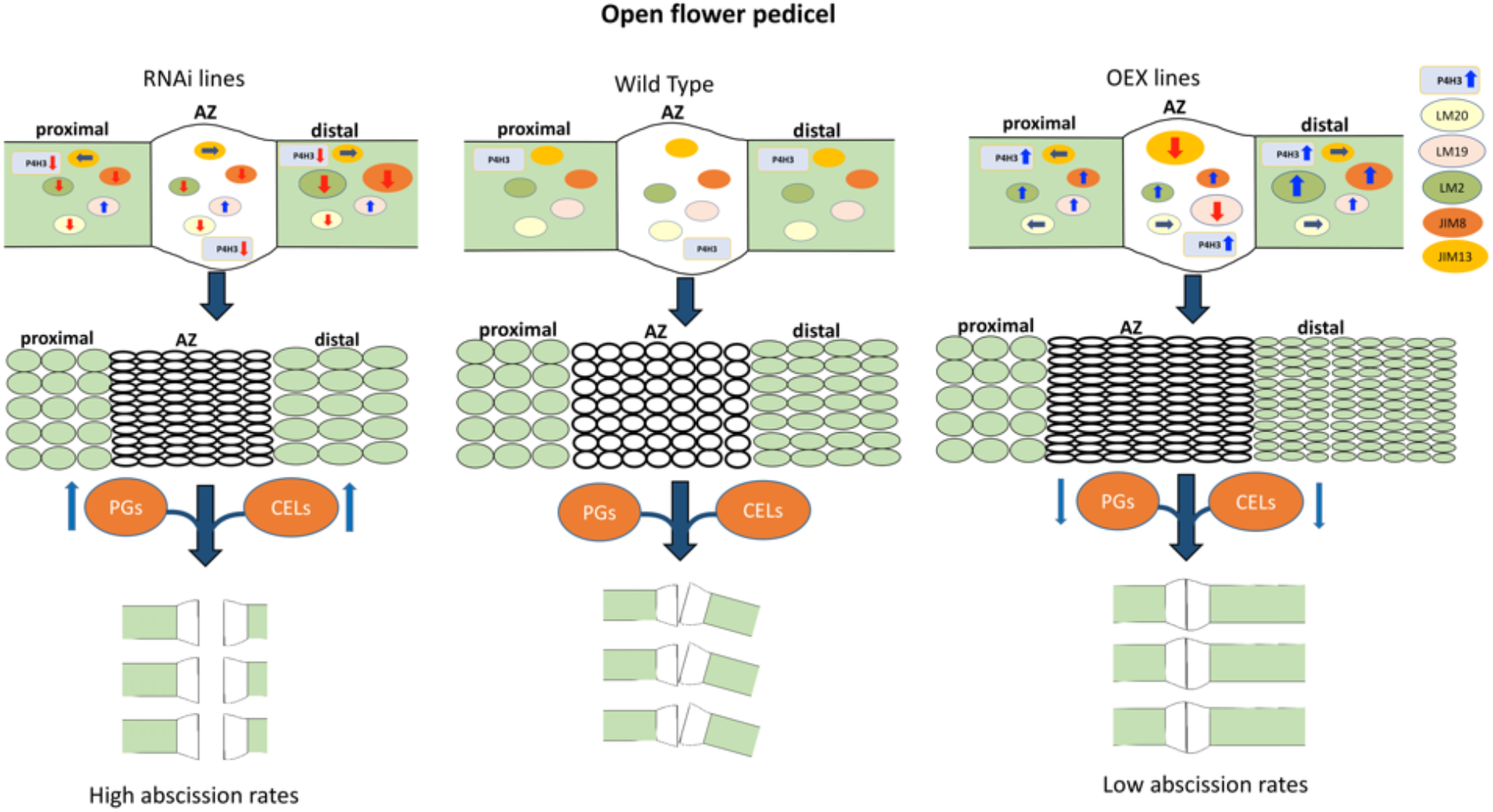
Schematic diagram depicting the molecular and morphological changes occurring in the open flower AZ and proximal and distal pedicel and their effect on ethylene-induced abscission rates of SlP4H3 OEX and RNAi lines. **A**. The SlP4H3 transcript is illustrated as rounded rectangular and the content of LM2, JIM8 and JIM13 AGP-bound epitopes and of highly methylesterified LM20-bound and partially demethyl-esterified LM19-bound HGs are illustrated as oval spheres of different colors in the AZ, proximal and distal pedicel of RNAi, OEX lines and wild type. The blue up-ward and the red down-ward arrows in the spheres indicate increase and decrease in content, respectively. The gray horizontal arrows indicate similar to wild type content. **B**. The morphological structure of the AZ and the pedicel is presented in relation to cell size, cell shape and pedicel length in the RNAi, OEX lines and wild type. There is a shift in the position of the AZ due to the length of the distal part of the pedicel. The distal part is longer and shorter in the OEX and RNAi lines, respectively. The number of AZ cell layers is higher in the OEX lines compared to wild type and RNAi lines as illustrated while the size and the shape of the AZ cells in both OEX and RNAi lines is smaller and more elongated. The cell size of the RNAi and OEX lines is larger and smaller compared to wild type, respectively. The oval spheres of PGs and CELLs represent the abscission-specific polygalacturonases and cellulases and the up-ward blue arrow indicates up-regulation in expression while the down-ward red arrow indicate down-regulation in expression. **C**. The illustration of abscised pedicels indicates higher abscission rate while the non-abscised pedicels indicate lower abscission rate. The partially abscised pedicels indicate partially abscised pedicels.

The shift in AZ position and the acceleration of ethylene-induced fruit abscission in OEX and RNAi was accompanied by molecular and morphological events associated with proline hydroxylated glycoproteins. There is the paradigm of AGP21 peptide which regulates root hair cell fate through disruption of a brassinosteroids-induced cascade of events (Borassi et al., 2020). There is also the IDA peptide which regulates flower abscission in Arabidopsis and requires proline hydroxylation for optimum activity (Butenko et al., 2014). In tomato, silencing of an IDA-like gene, SlIDL6, delayed low light induced flower abscission while the application of the SlIDL6 peptide accelerated abscission (Li et al., 2021). However, it is not known yet whether the SlIDL6 is proline hydroxylated or not.

Moreover, a tomato peptide hormone, phytosulfokine, was shown to enhance drought-induced tomato flower abscission by upregulation of TAPG2 and TAPG4 (Reichardt et al., 2020). However, there are no data up till now on proline hydroxylation of PSK (Kaufmann and Sauter, 2019). At a structural level, the distal part of the pedicel was shorter in RNAi and longer in OEX while the proximal pedicel was unaffected. Exactly the opposite phenotype was observed after ectopic expression of SLMBP21 which resulted in shorter proximal parts due to smaller cells (Liu et al., 2014). However, the shorter distal pedicel in the RNAi might be attributed to alterations in cell division of parenchymatic cells. The longer distal part in OEX lines might have been the result of enhanced cell division. The TAPG4 ectopic expression only in the distal pedicel due to overexpression of SlP4H3 suggests that alterations in the expression of cell division regulator(s) might also occur just in the distal part.

The tomato proximal and distal pedicels are characterized by distinct identity at the transcriptome level which might explain the changes in length of only the distal and not the proximal pedicel (Nakano et al., 2013). Moreover, suppression of SlMBP21 expression abolished the development of the flower AZ in a dose dependent manner (Liu et al., 2014), while suppression and overexpression of SlP4H3 resulted in smaller and more rectangular AZ cells and higher number of AZ cell layers in the OEX lines, indicating that SlP4H3 might independently regulate the AZ development programme per se.

The cell shape of the AZs in both the RNAi and OEX lines was more rectangular compared to wild type, possibly due to changes in the profile of AGPs as demonstrated by immunolocalization. Changes in cell shape were also obtained in hypocotyls of transgenic Arabidopsis with lower AGPs content, demonstrating a role for type II AGPs in the cell shape regulation (Yoshimi et al., 2020). At a structural level, the detachment force required for non-ethylene treated red ripe fruit AZ separation to occur was higher in RNAi compared to wild type and OEX, while the %E was also higher, implying greater resistance for tissue detachment. The greater force for detachment might be explained by the higher content of demethyl-esterified pectins in the flower AZs of RNAi lines which allows more extensive interchain cross-linking involving calcium ions which enhances cell wall stiffness and rigidity (Bidhendi and Geitmann, 2016; Cosgrove, 2016). Demethylesterified pectins might be involved in various developmental processes, such as the perception and transduction of extrinsic cues in protodermal cells (Giannoutsou et al, 2020a) and wall expansion during guard cell morphogenesis and opening of the stomatal pore (Giannoutsou et al., 2020b). Demethylesterified pectins might play a role in the above processes and the interplay between methylesterification and demethylesterification, might also be a determinant of the force required for the completion of the flower or fruit detachment. An additional factor is the thinner diameter of the AZ swollen node in OEX; i.e. a lower breaking force is required to detach AZ separation cell layers of thinner diameter. The thinner AZ diameter might be related to lower auxin content in the pedicel since it has been reported that the use of NPA (N-1-naphthylphthalamic acid), an auxin transport inhibitor, caused thickening of the distal pedicel of tomato fruit due to higher accumulation of auxin (Pattison and Catala, 2012). However, no auxin could be quantified in the red ripe fruit AZs of RNAi and OEX lines (data not shown).

In RNAi lines the LM2 and JIM8 AGPs were down-regulated in the flower AZs, proximal and distal parts, while in OEX lines the signal intensity was higher throughout pedicel (**Figure 6**). Therefore, the levels of LM2 and JIM8 AGPs might regulate the length of the pedicel since higher and lower content is associated with longer and shorter distal part, respectively. However, the JIM13 AGPs exhibited lower fluorescent signal in the AZ of OEX lines which is the only abscission-specific change in AGP(s) content among the three epitopes examined. This down-regulation was accompanied by a delay in progression of flower abscission. Therefore, the AZ-specific lower signal intensity of JIM13 AGP(s) might be related to alterations in the glycan structure due to changes in the frequency of occurrence of proline hydroxylation as a result of higher P4H activity in OEX lines.

Moreover, stable tomato transformants expressing SlP4H3-GFP fusion proteins were used for Co-IP of flower AZs protein extracts indicating interaction with the LM2, JIM13 and JIM8 AGPs-bound epitopes. These data suggest that these AGPs epitopes were hydroxylated by SlP4H3; therefore differences in their content in the RNAi and OEX lines might be due to SlP4H3 hydroxylation activity and not the result of another indirect regulatory mechanism.

The glycosylation of AGPs is considered very important for their function (Showalter 2016, Borassi et al., 2020). A decrease in JIM13 AGPs was also observed in olive fruit AZs, but it was associated with fruit abscission progression (Para et al., 2020). However, such association is in accordance with the decrease in LM2 and JIM8 AGPs found in the accelerated abscission of RNAi lines. In RNAi lines, the lower content of methyl-esterified HGs and the higher of demethyl-esterified HGs might be correlated to an acceleration of ethylene-induced abscission rate. The higher petal shedding rate in rose (*Rosa hybrid*), due to silencing of two ERFs (Ethylene Response Factors), was also associated with a reduction in highly methylesterified HGs (Gao et al., 2019). Moreover, higher levels of demethyl-esterified HGs were observed in olive fruit abscission which is consistent with the HG composition in RNAi lines (Para et al., 2020). These data support the involvement of methyl- and demethyl-esterified HGs with the abscission rate in tomato flower, olive fruit and rose petals (Gao et al., 2019; Para et al., 2020).

The AGPs might cross link with HGs of the cell wall pectins in the tomato pedicel cells. As a result, alterations either in the content or the structure of AGPs, due to higher or lower proline hydroxylation events, might affect the levels of cross linking to HGs causing changes in their exposure to pectic enzymes such as pectin methylesterases.

Alternatively, the arabinogalactan polysaccharides (AGs) and specifically the β-linked glucuronic acids were shown to be involved in developmental programs such as seedling growth in Arabidopsis through their capacity to act as apoplastic calcium repositories (Lopez-Hernandez et al., 2020). Therefore, it is possible that alterations in the frequency of occurrence of proline hydroxylation might affect AGPs content or their proper O-glycosylation, causing changes in abscission through alterations in the binding and release of apoplastic calcium.

At a molecular level, the higher expression of cell wall hydrolases such as TAPG1, TAPG2, TAPG4, CEL2 and CEL5 in the OEX lines, compared to those of RNAi, in the separation zone resulted in the advancement of abscission (Kalaitzis et al., 1997; Meir et al., 2010; Sundaresan et al., 2016). This might be attributed to changes on proline hydroxylation levels in substrate proteins such as regulatory AGPs.

In summary, the results of the present study indicate that P4Hs may regulate structural and molecular features of the abscission zones by modifying the structure and thereby the function of regulatory AGPs. Identification of AGPs interacting with SlP4H3 will provide further insight into the mechanism of tomato abscission regulation and unravel mechanistic details on the exact role of the post-translational modifications in the AGPs structures.

## Conclusions

Alterations in the expression of a tomato prolyl 4 hydroxylase, SlP4H3, caused a relocation in the position of the AZ either proximal or distal from the subtending organ. Moreover, ethylene-induced flower abscission was accelerated in SlP4H3 RNAi lines and delayed in OEX lines; whereas exactly the opposite response was observed in the red ripe fruit AZs. These directly opposite abscission progression trends could be partly attributed to changes in the abscission-specific cell wall hydrolases. The pedicel AZ responded in exactly the opposite way in a different developmental context. Therefore molecular mechanisms which were identified to regulate tomato flower abscission might not play a similar role in fruit abscission.

## Materials and Methods

### Generation of binary vectors, agro-mediated transformation and transgenic plants

The SlP4H3 overexpression vector (P4H3-OEX) were generated using the Gateway cloning technology (Invitrogen, Thermo Fisher Scientific, Waltham, MA, USA). Specific primers were designed comprising the attb sites to amplify the open reading frame of SlP4H3 gene (Solyc02g083390.2) (**Table S1**). The amplified P4H3 cDNA was subcloned into the pDONR221 entry vector and the construct was recombined in the appropriate destination Gateway® vector. The pK7WG2D.1 destination vector was used for the generation of the P4H3-OEX lines (Karimi et al., 2002). The construct integrity was verified by sequencing. Agrobacterium tumefaciens (strain LBA4404) was used for the transformation of tomato Ailsa Craig and Microtom cultivars.

The pK7WG2 destination vectors (/https://gatewayvectors.vib.be/collection) bearing the 35S promoter and eGFP for a C-terminal and N-terminal fusions were used for the generation of stable 35S:SlP4H3:eGFP and 35S:eGFP:SlP4H3 Ailsa Craig transformant lines.

The integration and construct integrity in the transformed plants was confirmed by PCR using the appropriate sets of primers (**Table S1**) while the homozygous plants were selected by using a Taqman qPCR approach and segregation analysis (German et al., 2003).

The independent Microtom transformant lines were validated by PCR (one cycle at 95°C for 5 min, 35 cycles at 94°C for 1 min, 54°C for 30s, 72°C for 30s, and extension cycle at 72°C for 10 min). The homozygous lines were crossed with the Ailsa Craig cultivar.

The experiments were conducted in homozygous T2 and T3 generation plants of two overexpression lines (OEX#1 and OEX#2). The Ailsa Craig tomato plants were grown in the experimental greenhouse facilities of the Mediterranean Agronomic Institute of Chania. The production of the RNAi-P4H3 lines (RNAi#1, RNAi#6 and RNAi#7) was previously described (Perrakis et al., 2019).

### Ethylene treatment

Open flowers and red ripe fruits were collected from the experimental greenhouse of the Mediterranean Agronomic Institute of Chania (MAICh) and the abscission explants, after removal of the subtending organs, were placed in beakers with water within gas-tight glass jars and exposed to constant flow of 10 ppm of ethylene while the abscission rate was determined at specific time intervals. The open flower AZs were collected and frozen in liquid nitrogen for further analysis after 0h, 4h and 8h of ethylene while the red ripe fruit AZs were collected after 0h, 24h, 48h and 72h of ethylene treatment.

### RNA extraction, cDNA synthesis and Quantitative Real-Time PCR analysis

Total RNA was isolated from flower and fruit AZs WT, two SlP4H3 OEX lines (#1, #2) and three SlP4H3 RNAi lines (#1, #6 and #7) by using the TRI reagent (Thermo Fisher Scientific, Waltham, MA, USA). Reverse transcription of approximately 1 μg of total RNA was performed with Superscript II® Reverse Transcriptase (Thermo Fisher Scientific, Waltham, MA, USA) and oligodT12-18 primers (Biotools, Spain). Quantitative real time PCR analysis was performed by using the CFX Connect™ Real-Time PCR Detection System (Bio-Rad) with SYBR™ Select Master Mix (Thermo Fisher Scientific, Waltham, MA, USA) sand gene specific primers (**Table S1**). The thermal cycling conditions were 50 °C for 2 min, 95 °C for 10 min followed by 95 °C for 15 s, 60 °C for 30 s and 72 °C for 30 s for 40 cycles. The cDNA samples were normalized with Actin (**Table S1**) and the data were analyzed by using the 2-ΔΔCT method (Livak and Schmittgen 2001). At least three biological replicates were performed for each gene expression experiment.

### Co-Immunoprecipitation and Western blot analysis

Immunoprecipitation was performed using GFP-Trap®A reagent (Chromotek) in flower abscission zones from wild type, GFP-P4H3 and P4H3-GFP plants according to manufacturer’s instructions. Bound proteins were eluted from the beads by boiling in 2× SDS sample buffer and analyzed by Western blot using LM2, JIM13, JIM8 (PlantProbes) and GFP (Chromotek) antibodies. The eluted proteins from wild type, N-GFP-P4H3 and C-GFP-P4H3 lines were extracted according to (Woodson 1987) with some modifications (Fragostefanakis et al., 2014). The membranes incubated with the LM2, JIM13 and JIM8 primary antibodies (PlantProbes, Leeds, UK). The Actin (Agrisera AB, Vännäs, Sweden) was used as a loading control. The bound antibodies were labeled with horseradish peroxidase-conjugated secondary antibodies (Agrisera AB,Vännäs, Sweden) and detected with ECL (Millipore, MA, USA) detection system. The range of the signal of the three bound epitopes was calculated using the ChemiDoc XRS system. At least three biological replicates were analyzed.

### Section embedding and toluidine blue O staining

Small pieces of pedicel that included the AZ area were fixed in 2% (w/v) PFA and 0.1% (v/v) glutaraldehyde in PEM at 4° C for 1.5 h. The specimens were washed in the same buffer and dehydrated in a graded ethanol series (10–90%) diluted in distilled water and three times in absolute ethanol for 30 min (each step) at 0° C. The material was post-fixed with 0.25% (w/v) osmium tetroxide added to the 30% ethanol step for 2 h. The material was infiltrated with LR White (LRW) (Sigma) acrylic resin diluted in ethanol, in 10% steps to 100% (1 h in each), at 4° C and with pure resin overnight. The samples were embedded in gelatin capsules filled with LRW resin and polymerized at 60° C for 48 h. Finally, the samples were sectioned using an ULTROTOME III TYPE 8801A ultramicrotome (LKB, Stockhom, Sweden) equipped with a glass knife. Semi-thin sections (0.5–2 μm) were stained with 0.5% (*w/v*) toluidine blue O and observed by light microscopy.

### Immunolocalization of homogalacturonans and AGPs

The distribution of HGs with a high degree of esterification recognized by LM20 antibody (Verhertbruggen et al. 2009) and HGs with a low degree of esterification recognized by LM19 antibody (Verhertbruggen et al. 2009) has been studied. Furthermore, for the distribution of AGPs epitopes, LM2, JIM8 and JIM13 antibodies have been tested. LM20, LM19, LM2, JIM8 and JIM13 (PlantProbes, Leeds, UK) were used as primary antibodies and FITC–conjugated anti-rat IgG (Sigma) as secondary antibody in all cases. All antibodies were diluted 1:40 in PBS that contained 2% (w/v) BSA.

Semithin sections (approximately 2-3μm thick) of material embedded in LRW resin were transferred to glass slides and blocked with 5% (w/v) BSA in PBS for 5 h. After washing with PBS, for the specimens where immunolabeling for AGPs was performed, an additional step for the removal of aldehydes has been included, by adding the sections in 1% glycine for 1h. After washing with PBS, the primary antibody diluted 1:40 in PBS containing 2% (w/v) BSA was applied overnight. Following rinsing with PBS and blocking again with 2% (w/v) BSA in PBS, the sections were incubated for 1 h, at 37° C in FITC anti-rat IgG (Sigma) diluted 1:40 in PBS containing 2% (w/v) BSA. After rinsing with PBS, the sections were mounted with a mixture of glycerol/PBS (2:1 v/v) containing 0.5 % (w/v) p-phenylenediamine (anti-fade medium).

### Observation and photography

The specimens were examined with a Zeiss Axioplan microscope equipped with a UV source, a Differential Interference Contrast (DIC) optical system, and a Zeiss Axiocam MRc5 digital camera. The filter set used for the specimens’ observation was provided with exciter pass band filter 450– 490 nm and barrier pass band filter 515–565 nm.

### Determination of mechanical properties of tomato pedicels

Tensile tests were performed on tomato pedicels with a TA-XT2i Texture Analyzer (Stable Microsystems, Godalming, Surrey, UK) at 25°C using tensile grips (A/TG) to evaluate the mechanical properties of the pedicel abscission zone of the fruit. The tomato pedicel, including the abscission zone, were mounted on the grips and a tensile test with deformation up to break at a crosshead speed of 0.8 mm/s was applied. The pedicels were always broken at the abscission zone. At least twenty samples for each tomato line were measured. The force–displacement curves were used to obtain the abscission force of the fruit (F_max_) which was the force recorded at knuckle fracture. From these curves, the Young’s modulus (E) can be calculated from the initial slope of the stress–strain curves, the maximum force at break as well as the percentage elongation of tomato pedicel (%*E*) at fracture using the following equations: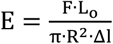 and 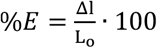, where R is the radius of pedicle knuckle, F is the force at the initial slope of the curve, Δl and ΔL is the increase of pedicel length at the initial slope of the curve and at failure, respectively and L_o_ is the gauge length (i.e. the initial length of the test section).

### Cell Parameter Measurements

To quantify the differences in the morphology of AZ cells of flower and fruits AZs WT, two SlP4H3 OEX lines (#1, #2) and three SlP4H3 RNAi lines (#1, #6 and #7) we took measurements of cell width, height, section area, and perimeter from a series of images. The slides were edited in ImageJ to carry out image related routine operations, such as color thresholding and amplification (Abràmoff et al., 2003). Cell parameter measurements were performed in a selected area within each slide from the microscopic images for calculation of cell width, cell area and other cell parameters by using the ImageJ program.

### Statistical Analysis

All measurements were conducted in triplicate and results are re-express as mean ± standard errors (SEs). Then, all the data were subjected to statistical analysis of variance (ANOVA) and a *post hoc* multiple comparison test (Tukey’s honest significant difference criterion, 95% confidence interval) by using the statistics toolbox of MATLAB (The Mathworks Inc., Natick, MA, USA). Comparisons were made to identify any statistically significant differences between wild type (WT) and each RNAi and OEX line, respectively; differences at p < 0.05 were considered to be significant.

## Acknowledgments

We would like to thank Dr Mark Tucker for providing the seeds of TAPG4:GUS line. This work has been supported partly by the project “Tomato P4Hs” (Ref. Number 4388), which was implemented by “ARISTEIA II,” an Action of the “Operational Programme Education and Life Long Learning,” co-funded by the European Social Fund (ESG) and National Resources. Moreover, this research was partially supported by the project “PlantUp: Upgrading Plant Capital” (MIS 5002803) which is implemented under the Action “Reinforcement of the Research and Innovation Infrastructure,” funded by the “Operational Programme, Competitveness, Entrepreneurship and Innovation” (NSRF 2014-2020) and co-financed by Greece and European Union (European Regional Development Fund). Finally, this research has been partially co-financed by the European Union and Greek national funds through the Operational Program Competitiveness, Entrepreneurship and Innovation, under the call RESEARCH – CREATE – INNOVATE (project code: T2EDK-01332: TOMATOMICS-Development Of New Tomato Cultivars By Using Omics Technologies, MIS 5072532).

---

**Supplementary Figure 1.**
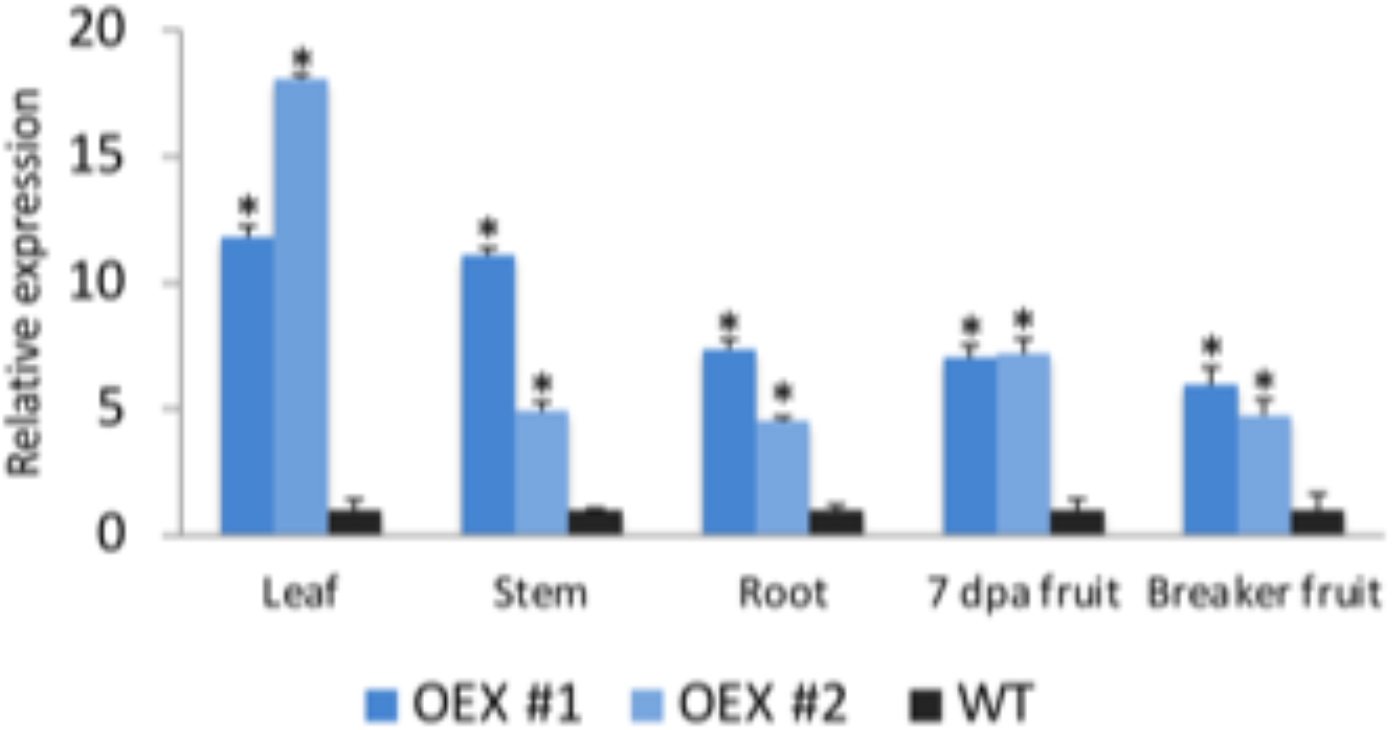
Tissue-specific expression of SlP4H3 in the OEX#1, OEX#2 lines. Relative expression of SlP4H3 in leaf, stem, root, 7 dpa (days post anthesis) fruit and Breaker stage fruit.

**Supplementary Figure 2.**
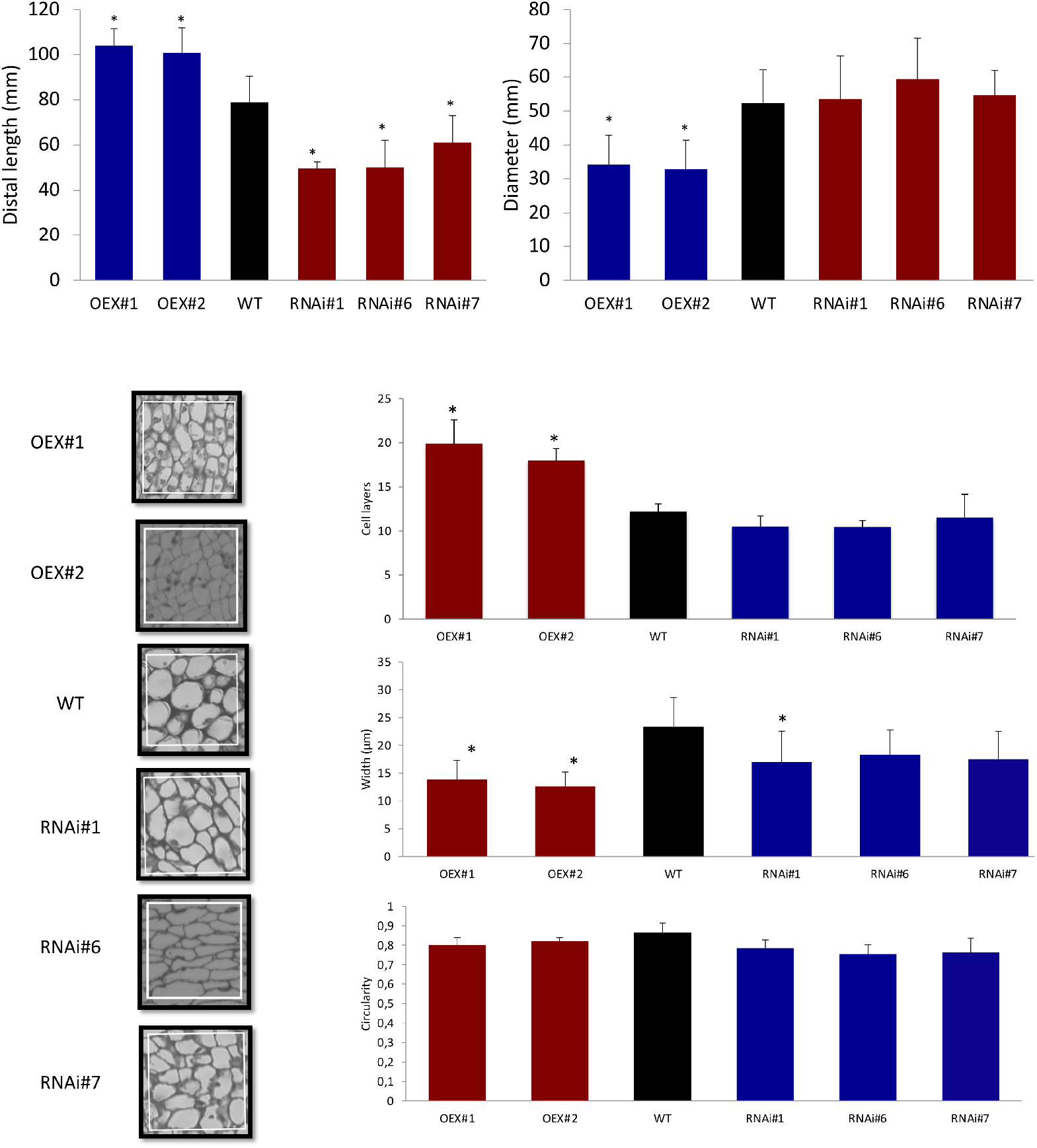
Red ride fruit Pedicel characterization in the RNAi #1, #6, #7 and OEX #1, #2 lines and wild type. **A**. The length of the distal part of the pedicel and the AZ diameter of RNAi #1, #6, #7 and OEX #1, #2 lines and wild type (n=180 pedicels/line). **B**. Longitudinal sections depicting indicative cell morphologies of distal pedicel parenchymatic cells in a specific square area of length 82μm. **C. Number of AZ cell layers**. The AZ cell layers in both OEX lines appear to be significant more compared to WT and RNAi lines. **Cell width**. The width/length of both the OEX AZ cells appeared to be significantly lower than WT. **Cell circularity**. The shape in the OEX and RNAi lines appeared to be more rectangular in comparison to the WT, in which they have higher circularity value. Cell morphology measurements made on 9 images per line; three regions per AZ from three different plants of the same line. The asterisks indicates statistically significant differences compared to wild type.

**Supplementary Table 1.**
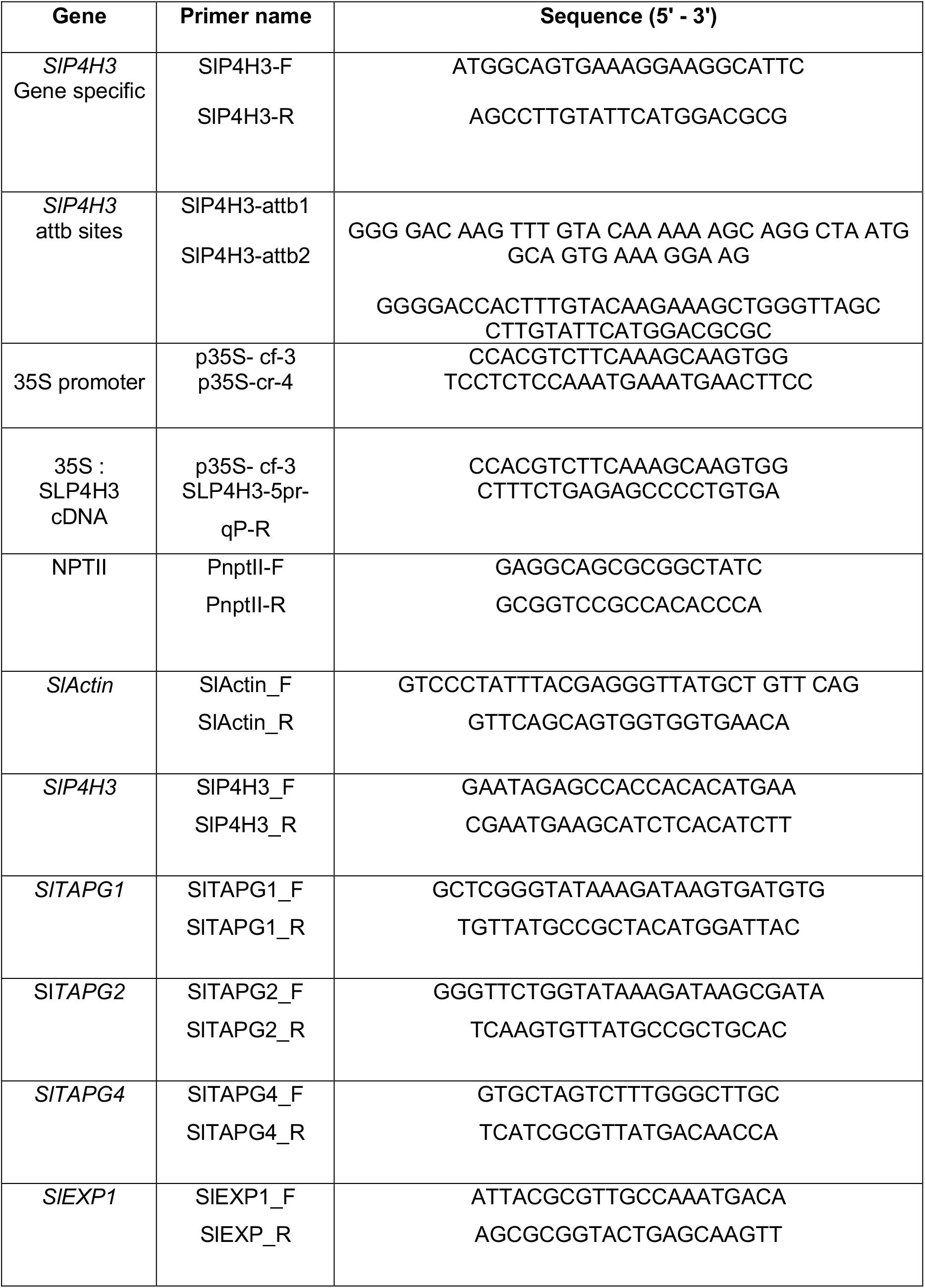
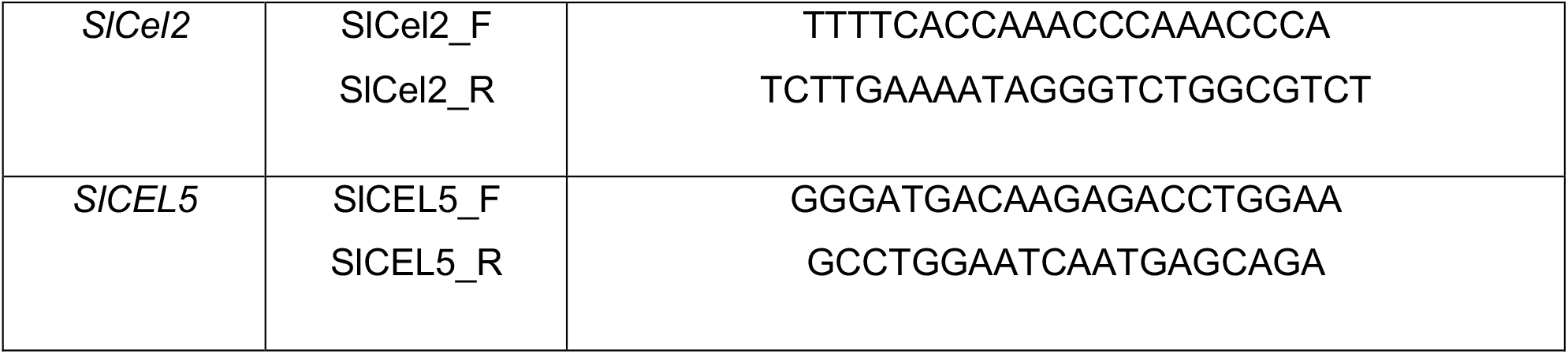
Table of primers used for the qPCR analysis, the subcloning of SlP4H3 cDNA in GATEWAY vector, and amplification of NPTII partial cDNA fragment.

